# Response diversity can stabilize or destabilize community dynamics depending on the number of insensitive species

**DOI:** 10.64898/2026.08.11.743952

**Authors:** Shota Shibasaki, Hiroaki Fujita, Hirokazu Toju, Masato Yamamichi

## Abstract

Investigating the factors that stabilize biological communities is a central topic in ecology. Response diversity, defined as variation in species responses to environmental change, has been proposed as a key mechanism underlying the biodiversity-ecosystem functional stability (BEFS) relationship, whereby greater species diversity enhances ecological stability. Previous studies have shown that response diversity promotes ecological stability by generating asynchronous population fluctuations and the resulting compensatory dynamics. Although several metrics have been proposed to quantify response diversity, they do not explicitly consider the presence of *insensitive* species whose performance is unaffected by current environmental conditions. To examine how insensitive species influence response diversity, species persistence, and ecological stability, we conducted numerical simulations of a generalized Lotka–Volterra model under environmental forcing. We first confirmed that increasing variation among sensitive species increased the response diversity index and stabilized community dynamics. We then examined a scenario in which response diversity depended solely on the proportion of sensitive and insensitive species, assuming that all sensitive species responded identically to environmental change. Under this assumption, the response diversity index was maximized when sensitive and insensitive species occurred in equal proportions, whereas increasing the number of sensitive species monotonically destabilized community dynamics. Consequently, the relationship between response diversity and community stability depended on how response diversity was generated, such that higher response diversity could even be associated with lower community stability. These findings demonstrate that overlooking environmentally insensitive species can obscure the mechanisms linking response diversity and ecological stability. More broadly, our results reveal that response diversity comprises at least two distinct biological components—species sensitivity and response variation among sensitive species—that can have contrasting consequences for community stability. We therefore highlight the need to quantify sensitive species empirically and to develop response diversity metrics that distinguish these components.

**Author Summary:** Understanding why some communities remain stable despite environmental change is a longstanding goal in ecology. Response diversity, which refers to differences in how species respond to environmental change, has been proposed as a key mechanism explaining why greater biodiversity (species richness) can promote ecological stability. Because species respond differently to changing environments, declines in some species can be compensated by increases in others, helping to stabilize community dynamics. However, previous studies have rarely considered species that are insensitive to current environmental changes. Using a mathematical model, we show that response diversity can arise from two distinct biological components—the number of sensitive species and variation in their responses—and that these components can have contrasting effects on ecological stability. When response diversity reflects variation among sensitive species, greater response diversity stabilizes community dynamics, as expected. In contrast, when response diversity changes only because of the proportions of sensitive and insensitive species, higher response diversity can be associated with lower community stability. Our findings highlight the importance of quantifying the number of sensitive species and developing response diversity metrics that distinguish species sensitivity from variation in responses among sensitive species.

## 1 Introduction

Understanding the factors that stabilize biological communities and ecosystem functioning has long been a central goal in ecology. Numerous studies have shown that greater species richness enhances ecosystem functioning and its temporal stability (Tilman et al. 1996, Isbell et al. 2009, Brun et al. 2019, Wang et al. 2024). In particular, biodiversity-ecosystem functional stability (BEFS) relationships have become a central concept in ecology because they provide a framework for understanding how biodiversity contributes to stable ecosystem functioning. Several theoretical frameworks have been proposed to explain BEFS relationships, including the insurance hypothesis (Yachi and Loreau 1999, Loreau et al. 2021) and the portfolio effect (Doak et al. 1998, Thibaut and Connolly 2013), both of which emphasize that greater species richness can enhance the temporal stability of ecosystem functioning. However, understanding the mechanisms underlying BEFS relationships remains a major challenge in ecology.

Among the mechanisms proposed to explain BEFS relationships, response diversity (Elmqvist et al. 2003, Mori et al. 2013) has emerged as a key mechanism linking biodiversity to ecological stability (Ives et al. 1999, Yachi and Loreau 1999, Hooper et al. 2005, Winfree and Kremen 2009, Bartomeus et al. 2013, Sasaki et al. 2019, Ross et al. 2023, Ross and Sasaki 2024, Danet et al. 2025, Hsieh et al. 2026, Ross et al. 2026). Response diversity is defined as variation in how species contributing to the same ecosystem function respond to environmental fluctuations (Laliberté et al. 2010). High response diversity is expected to stabilize ecosystem functioning because declines in some species can be compensated by increases in others. Previous theoretical and empirical studies suggest that differences in species’ environmental optima can generate asynchronous population dynamics, thereby stabilizing ecosystem functioning (Sasaki et al. 2019, Muraina et al. 2021, Schnabel et al. 2021, White et al. 2023). Accordingly, identifying the biological processes that generate response diversity is essential for understanding the mechanisms underlying BEFS relationships.

However, one fundamental aspect of response diversity has received little attention: the number of sensitive and insensitive species within a community. If most species in a community are insensitive to environmental change, overall response diversity remains low because variation is confined to only a small subset of species, even when sensitive species differ greatly in their responses. The presence of insensitive species does not imply that the environmental variable is ecologically unimportant for those species. For example, species with broader environmental response curves than others may appear insensitive if current environmental fluctuations fall within the flat portion of their response curves. Empirical studies have reported the coexistence of sensitive and insensitive species in natural communities (Fründ et al. 2013, Matsumae et al. 2025), yet their contribution to response diversity has rarely been examined explicitly. These observations suggest that similar levels of response diversity may arise through different biological mechanisms, potentially leading to contrasting consequences for ecological stability.

Here, we investigated how the number of sensitive species and variation in their responses influence response diversity, ecological stability, and species persistence using a mathematical model with randomly generated species interactions (May 1972). We found that the relationship between the response diversity index and community stability depends on how response diversity is generated, specifically through changes in the number of sensitive species or variation in their responses. As expected, increasing variation among sensitive species increased the response diversity index and stabilized community dynamics. In contrast, when response diversity changed solely through changes in the number of sensitive species, higher response diversity could be associated with lower community stability. Our findings highlight the importance of quantifying the number of sensitive species in natural communities and developing response diversity metrics that separately capture species sensitivity and variation in responses among sensitive species.

## 2 Model

### 2.1 Community dynamics

We modeled the dynamics of an *N*-species community under environmental fluctuations (Fig. 1A) using a generalized Lotka–Volterra (gLV) model, following previous studies (May 1972, Akjouj et al. 2024). The community dynamics are described by:

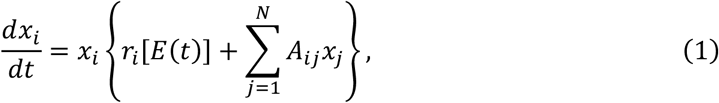

**Figure 1.**
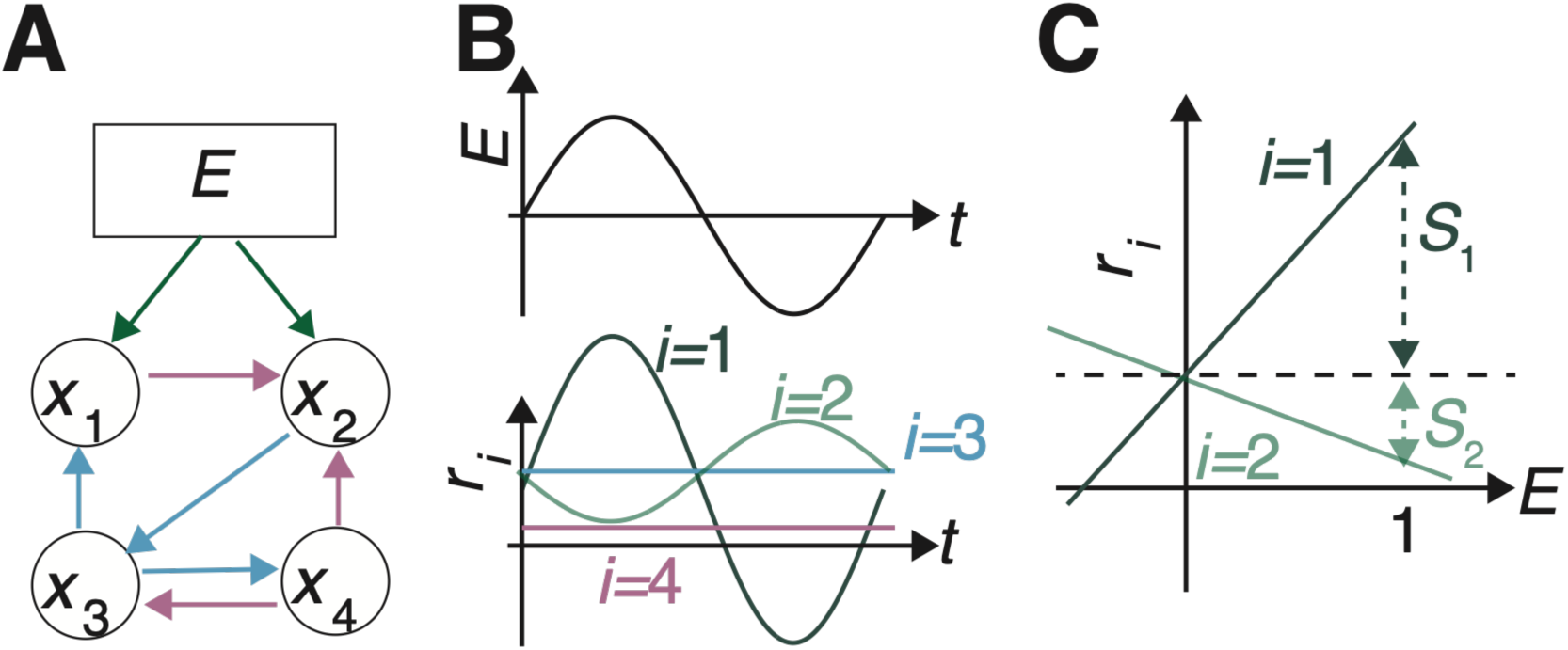
Overview of the theoretical framework. (A) Example of a four-species community under environmental fluctuations. Positive (blue) and negative (purple) arrows indicate species interactions. Species 1 and 2 are sensitive to the environmental variable *E* (green arrows), whereas species 3 and 4 are insensitive to the environmental variation considered here (no arrows from *E*). (B) Dynamics of the environmental variable *E*(*t*) (top) and corresponding intrinsic growth rates *r_i_*[*E*(*t*)] of the four species (bottom). Environmental sensitivity (*s_i_*) determines the amplitude of fluctuations in intrinsic growth rate; in this example, *s*_1_ > *s*_2_ > *s*_3_ = *s*_4_ = 0. Species 1 and 2 have different environmental optima (*γ*_1_ = 0 and *γ*_2_ = π), whereas species 3 and 4 maintain constant growth rates because they are insensitive to environmental fluctuations considered. (C) Plotting intrinsic growth rates against the environmental variable yields the slope *S_i_*. The variation in slopes among species forms the basis for quantifying response diversity.

where *x_i_* is the abundance of species *i*, *r_i_*[*E*(*t*)] is the intrinsic growth rate of species *i* as a function of the environment *E*(*t*), which is defined using a sine function (see *Section 2.2*), and **A** = {*A_ij_*} is the species interaction matrix.

We fixed the intraspecific interaction coefficients at *A_ii_* = –1 for all *i* = 1, …, *N*, so that *r_i_*[*E*(*t*)] corresponds to the single-species equilibrium abundance (i.e., carrying capacity) of species *i* under environment *E*(*t*). For interspecific interactions (*i* ≠ *j*), each interaction was present with probability *C* (connectance). We considered *C* = 0, 0.1, 0.25, 0.5, 0.75, and 1. Interaction coefficients were drawn from a normal distribution with mean 0 and standard deviation (SD) 0.2. This SD was chosen to reduce the occurrence of unrealistically unstable dynamics caused by excessively strong positive interactions. Thus, for *i* ≠ *j*, *A_ij_* ∼ *N*(0, 0.2^2^) with probability *C*, whereas *A_ij_* = 0 with probability 1 – *C*.

We ran simulations with *N* = 100 until *t* = 5,000. Initial species abundances were set to *x_i_*(0) = 0.1 for all species. We generated 1,000 species interaction matrices and examined the effects of connectance (*C*) and environmental fluctuations on community dynamics. We performed simulations using the ode function with the fourth-order Runge-Kutta (rk4) method and a step size of 0.1 in the deSolve package version 1.38 (Soetaert et al. 2010) in R version 4.3.1 (R Core Team 2024). We set the extinction threshold to *x_i_*(*t*) < 0.001. Once a species fell below this threshold, its abundance was set to zero thereafter.

At the end of each simulation, we evaluated two community properties: realized species richness (i.e., the number of species whose abundance exceeded the extinction threshold of 0.001) and the coefficient of variation (CV) of total community abundance (hereafter, abundance CV), which served as a proxy for community instability (Leary and Petchey 2009, Hordley et al. 2021). The abundance CV was calculated as the ratio of the SD to the mean of total community abundance using the final 100 time units (over the interval from *t* = 4,900 to *t* = 5,000) to evaluate temporal variability after transient dynamics had dissipated.

Some simulations terminated before *t* = 5,000 because one or more species abundances increased without bound. These simulations were excluded from subsequent analyses. Such excluded simulations occurred more frequently at higher connectance values (see *Section 3*). When all species went extinct, the abundance CV was set to zero because total community abundance remained constant at zero.

### 2.2 Implementation of environmental fluctuations

We assumed that the environmental variable *E* ranged between –1 and 1. Specifically, we modeled environment fluctuations as a sinusoidal function over time, *E*(*t*) = sin(*βt*), and assumed that the environmental variable affected the intrinsic growth rates of species *r_i_*[*E*(*t*)] (Fig. 1B). This type of environmental variation represents, for example, seasonal changes in temperature. Although environmental conditions can also alter species interactions (Chamberlain et al. 2014, Piccardi et al. 2019, Meacock and Mitri 2025), we assumed that the environmental variable affected only intrinsic growth rates because previous studies have quantified response diversity based on variation in intrinsic growth rates across environmental conditions (Leary and Petchey 2009, McCann 2016).

The intrinsic growth rate of species *i* under environmental fluctuations was defined as:

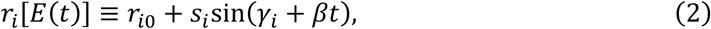

where *r_i_*_0_ is the baseline intrinsic growth rate of species *i*, *s_i_* is the sensitivity to environmental fluctuations, *γ_i_* takes a value of 0 or π and determines whether species *i* attains its maximum growth rate at *E* = 1 or *E* = –1, and *β* determines the period of environmental fluctuations. Baseline growth rates were sampled from a uniform distribution, *U*(0, 1). We fixed *β* = 0.04, which generated environmental fluctuations at an intermediate rate. Under slower fluctuations, environmental change had little effect because communities approached equilibrium before environment changed substantially, whereas under faster fluctuations the environment effectively averaged over time (Shibasaki et al. 2021). The sensitivity parameter *s_i_* and the phase parameter *γ_i_* varied among the simulation scenarios described below. Table 1 summarizes the model variables and parameters.

**Table 1:** A list of variables and parameters.

| Symbol | Description | Value or distribution |
| --- | --- | --- |
| $x_i$ | Abundance of species $i$ | $\geq 0$ |
| $N$ | Number of species in a community | 100 |
| $A_{ij}$ | Species interaction coefficient | $-1$ for $i = j$ and $N(0, 0.2^2)$ with the probability $C$ and otherwise $0$ for $i \neq j$ |
| $r_{i0}$ | Baseline intrinsic growth rate of species $i$ | $U(0, 1)$ |
| $s_i$ | Species $i$'s sensitivity to the environment | $0$ or $U(0, 1)$ |
| $\gamma_i$ | Phase of growth rate fluctuations | $0$ or $\pi$ |
| $\beta_i$ | Periods of environmental changes | 0.04 |

$U(a, b)$ represents uniform distribution between $a$ and $b$ . $N(a, b^2)$ represents normal distribution
whose mean is $a$ and standard deviation is $b$ .

To disentangle the effects of the number of sensitive species and variation in their responses on response diversity and community dynamics, we analyzed three scenarios that differed in species’ environmental responses.

#### 2.2.1 Scenario 1: No Response to the Environmental Fluctuations

We first considered a baseline scenario in which no species responded to environmental fluctuations (*s_i_* = 0 for *i* = 1, …, *N*). This scenario is equivalent to those examined in classic studies of the complexity–stability relationship in ecological communities (Gardner and Ashby 1970, May 1972, Allesina and Tang 2012). It served as a baseline for evaluating realized species richness and abundance CV when no species responded to environmental fluctuations. For this scenario, we generated 1,000 interaction matrices for each of the six connectance values, resulting in 6,000 simulations in total.

#### 2.2.2 Scenario 2: Binary Variation in Species’ Responses

We next considered a scenario in which response diversity varied solely through changes in the number of sensitive species, while all sensitive species shared an identical environmental response. We designated *M* species (*i* = 1, …, *M*) as sensitive and the remaining *N* – *M* species (*i* = *M* + 1, …, *N*) as insensitive, where *M* ≤ *N*. We considered *M* = 1, 5, 10, 25, 50, and 100. Sensitive species were assigned *s_i_* = 0.5 and *γ_i_* = 0, whereas insensitive species were assigned *s_i_* = 0 and *γ_i_* = 0. We chose *s_i_* = 0.5 because it corresponds to the mean sensitivity in Scenario 3. For each combination of connectance and *M*, we generated 1,000 interaction matrices, resulting in 36,000 simulations in total.

#### 2.2.3 Scenario 3: Quantitative Variation in Species’ Responses

We extended Scenario 2 by allowing quantitative variation in the responses of sensitive species. For the *M* sensitive species (*i* = 1, …, *M*), the sensitivity parameters (*s_i_*) were independently sampled from a uniform distribution, *U*(0, 1). This distribution has a mean of 0.5, matching the fixed sensitivity used in Scenario 2 while introducing variation among sensitive species. *γ_i_* was assigned a value of 0 or π with equal probability. The values of *M* and the parameter settings for insensitive species were identical to those in Scenario 2. For each combination of connectance and *M*, we generated 1,000 interaction matrices, resulting in 36,000 simulations in total.

### 2.3 Measuring response diversity

Following Ross et al. (2023), we quantified response diversity based on variation in the slope of the intrinsic growth rate with respect to the environmental variable *E*. The slope of species *i*’s response, *S_i_*, was defined as (Fig. 1C):

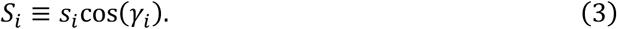

Equation (3) shows that *s_i_* determines the magnitude of the slope, whereas *γ_i_* determines its sign.

Ross et al. (2023) introduced two response diversity metrics: dissimilarity and divergence. The dissimilarity metric (hereafter, response dissimilarity) is a special case of the diversity of order 0 defined by Leinster and Cobbold (2012):

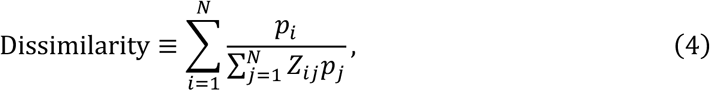

where *p_i_* is the relative abundance of species *i*, and *Z_ij_* is the similarity between the environmental response slopes of species *i* and *j*:

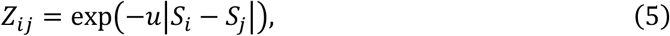

where *u* is a positive constant. In our simulations, *p_i_* was calculated as the mean relative abundance of species *i* over the interval from *t* = 4,900 to *t* = 5,000, and we fixed *u* = 1. The choice of *u* did not qualitatively affect the results. We defined the response dissimilarity as 1 when all species went extinct or when only a single species remained, because this is the minimum possible value of the metric.

Ross et al. (2023) also defined the divergence metric of response diversity (hereafter, response divergence) based on the difference between the maximum and minimum response slopes. Because the intrinsic growth rate is linear with respect to the environmental variable *E*, the divergence metric simplifies to:

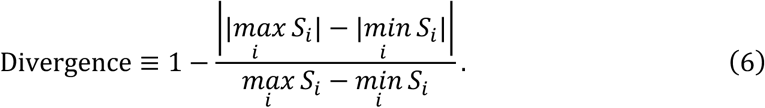

Response divergence is positive only when 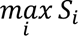 and 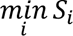 have opposite signs; otherwise, it equals zero. Its maximum value is 1, which occurs when 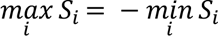.

We focused on response dissimilarity in the following analyses because response divergence was effectively binary in our simulations. In Scenarios 1 and 2, response divergence was always zero by definition. In Scenario 3, it was either close to 0 or close to 1 (Fig. S1A). Therefore, response divergence was not informative for analyzing relationships with realized species richness or abundance CV. In contrast, response dissimilarity increased continuously with the number of sensitive species, although there was some scatter around this trend (Fig. S1B).

### 2.4 Statistical analysis

All statistical analyses were performed in R version 4.3.1 (R Core Team 2024). We used quantile logistic regression to analyze realized species richness and quantile linear regression to analyze abundance CV. Quantile logistic regression was performed using the Log.lqr() function in the lqr package (version 5.0; Galarza et al. 2021). Quantile linear regression was performed using the rq() function in the quantreg package (version 5.97; Koenker 2025). Akaike’s information criterion (AIC) was calculated using the aic() function.

## 3 Results

### 3.1 Scenario 1: Higher connectance decreased species richness but did not affect ecological stability

We first examined Scenario 1, in which no species responded to environmental fluctuations. We excluded 460 of the 6,000 simulations because one or more species abundances became unbounded, mostly at high connectance values (2 simulations at *C* = 0.5, 101 at *C* = 0.75, and 357 at *C* = 1). Consistent with May (1972), increasing connectance reduced realized species richness (Fig. S2A). In contrast, abundance CV remained low across connectance values (Fig. S2B), suggesting that most simulations converged to equilibrium. These results provide a baseline for the following analyses.

### 3.2 Scenario 2: The number of sensitive species generated contrasting relationships between response diversity and stability

We next examined Scenario 2, in which species were either sensitive or insensitive to the environmental variable. We varied the number of sensitive species while assuming that all sensitive species responded identically to environmental change. Of the 36,000 simulations, 33,265 were retained for analysis; excluded simulations occurred primarily at high connectance values (8 at *C* = 0.5, 555 at *C* = 0.75, and 2,172 at *C* = 1). Realized species richness decreased with increasing connectance and the number of sensitive species, although these variables showed a significant positive interaction (Fig. 2A; Table S1). In contrast, abundance CV was largely independent of connectance but increased with the number of sensitive species (Fig. 2B; Table S2).

**Figure 2.**
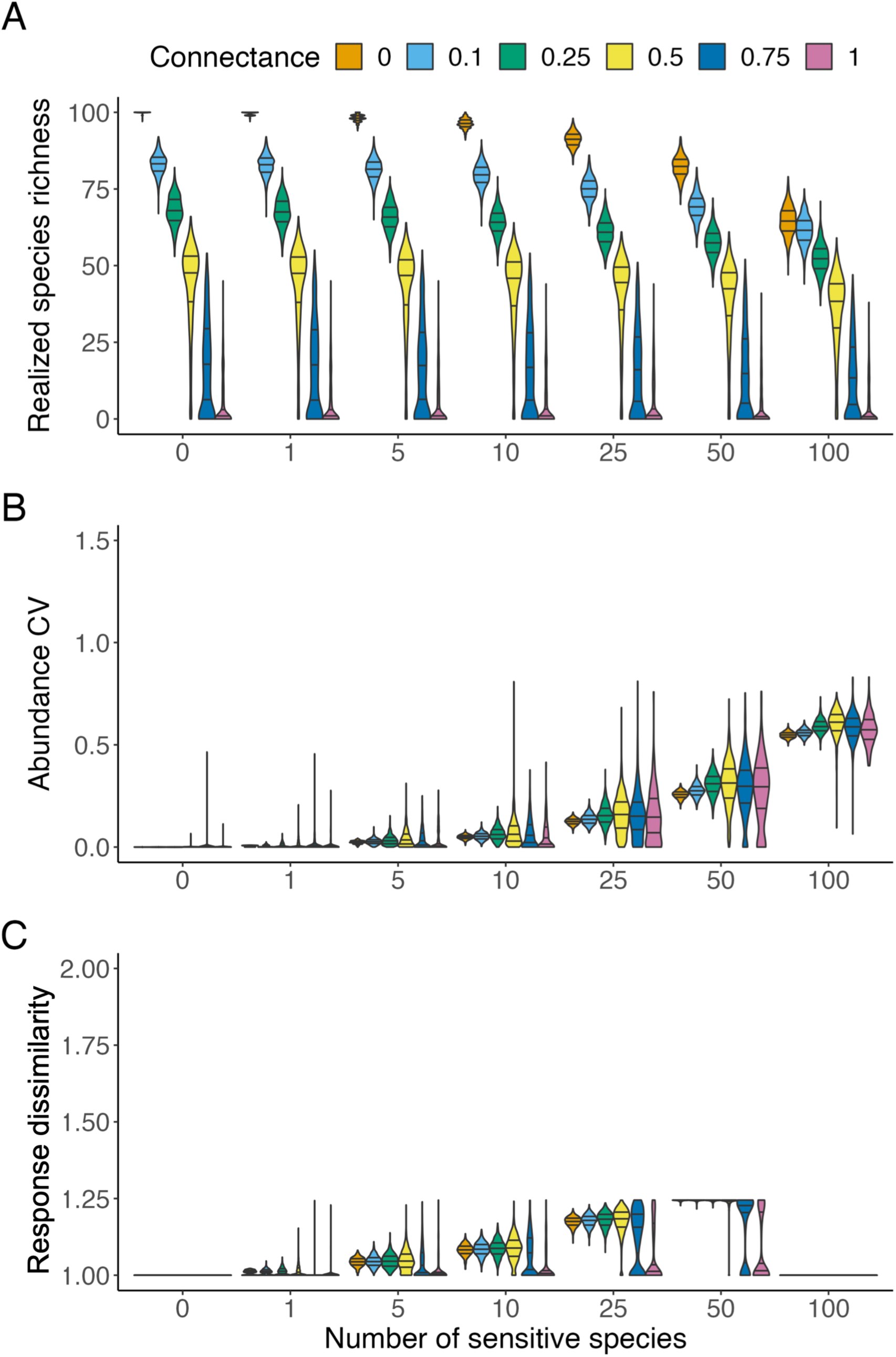
Realized species richness, abundance CV, and response diversity under binary variation in species’ responses to environmental change. Realized species richness (A), abundance CV (B), and the dissimilarity metric of response diversity (response dissimilarity; C) are shown as functions of the number of sensitive species and connectance. Horizontal lines indicate fitted quantile regression estimates. Note that response dissimilarity showed a bimodal distribution when the number of sensitive species was 25 or 50 and connectance was high (0.75 or 1). This occurred because response dissimilarity was set to one when zero or one species persisted, whereas it was generally higher when two or more species persisted.

Whereas realized species richness and abundance CV changed monotonically with the number of sensitive species, response dissimilarity showed a unimodal relationship with the number of sensitive species (Fig. 2C). This pattern follows directly from the properties of the response dissimilarity metric (Leinster and Cobbold 2012), which can be interpreted as a similarity-sensitive version of species richness. When all species had the same sensitivity (all sensitive or all insensitive to the environmental variable), response dissimilarity took its minimum value of 1 because there was no variation among species’ responses. In contrast, response dissimilarity was maximized when half of the species were sensitive. Because response dissimilarity changed non-monotonically with the number of sensitive species, it showed a poorer fit than the number of sensitive species for predicting abundance CV (higher AIC; Table S2). However, response dissimilarity showed a better fit for predicting realized species richness than the number of sensitive species (Table S1).

### 3.3 Scenario 3: Quantitative variation in species’ responses weakens the destabilizing effect of sensitive species

In Scenario 3, we allowed quantitative variation in species’ environmental responses such that sensitive species differed in the slopes of their environmental responses. Of the 36,000 simulations, 33,210 were retained for analysis. Excluded simulations occurred primarily at high connectance values (11 at *C* = 0.5, 588 at *C* = 0.75, and 2,191 at *C* = 1). The pattern of realized species richness (Fig. 3A) was similar to that observed in Scenario 2 (Fig. 2A). In contrast, abundance CV (Fig. 3B) and response dissimilarity (Fig. 3C) showed different patterns from those in Scenario 2 (see also Figs. S3 and S4). Abundance CV was lower than in Scenario 2 (Fig. 2B), even when all species were sensitive to the environmental variable. Quantile regression further showed that the effect size of the number of sensitive species on abundance CV was approximately tenfold smaller in Scenario 3 (Table S3) than in Scenario 2 (Table S2). Unlike Scenario 2, response dissimilarity increased monotonically with the number of sensitive species (Fig. 3C). Consequently, using response dissimilarity instead of the number of sensitive species improved the model fit for both realized species richness (Tables S3) and abundance CV (Table S4).

**Figure 3.**
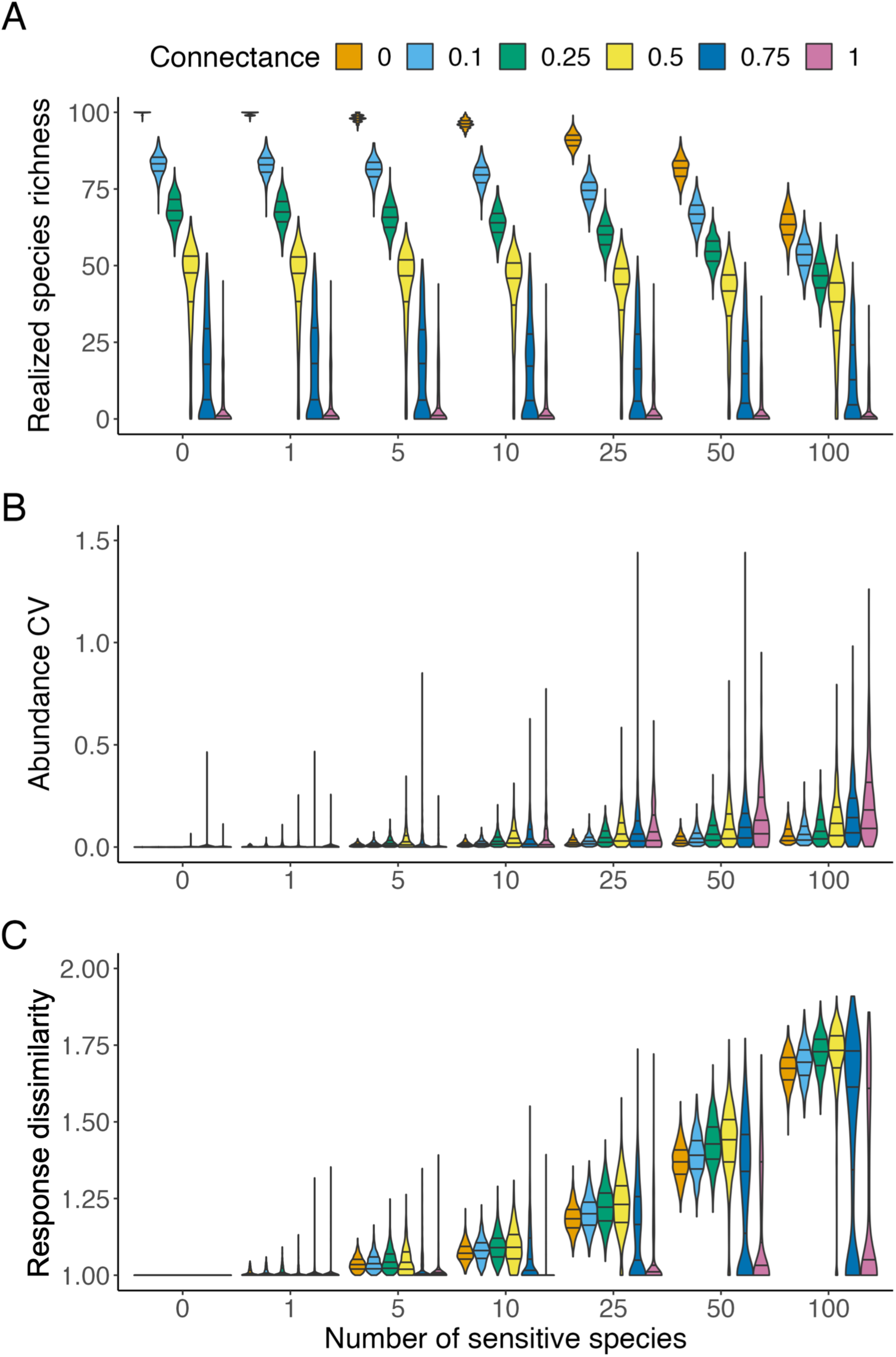
Realized species richness, abundance CV, and response diversity under quantitative variation in species’ responses to environmental change. Realized species richness (A), abundance CV (B), and response dissimilarity (C) are shown as functions of the number of sensitive species and connectance. This figure presents the results of Scenario 3, in which sensitive species differed in the magnitude and direction of their responses to their environmental response slopes.

### 3.4 The number of sensitive species altered the relationship between response diversity and stability

Comparing Scenarios 2 and 3 revealed how response dissimilarity, abundance CV, and the relationship between them changed with the number of sensitive species and variation among their responses. Figure 4 illustrates representative examples with connectance fixed at *C* = 0.5. When variation among sensitive species was introduced while the number of sensitive species was held constant, abundance CV decreased with increasing response dissimilarity (yellow arrow in Fig. 4). This negative relationship between response dissimilarity and abundance CV (i.e., the stabilizing effect of response diversity) is consistent with the expectation of previous studies. In contrast, when the number of sensitive species varied, abundance CV either increased (green arrow in Fig. 4) or decreased (blue arrow in Fig. 4) with response dissimilarity. This pattern arose because response dissimilarity was maximized at intermediate numbers of sensitive species (Fig. 2C), whereas abundance CV increased monotonically with the number of sensitive species (Fig. 2B).

**Figure 4.**
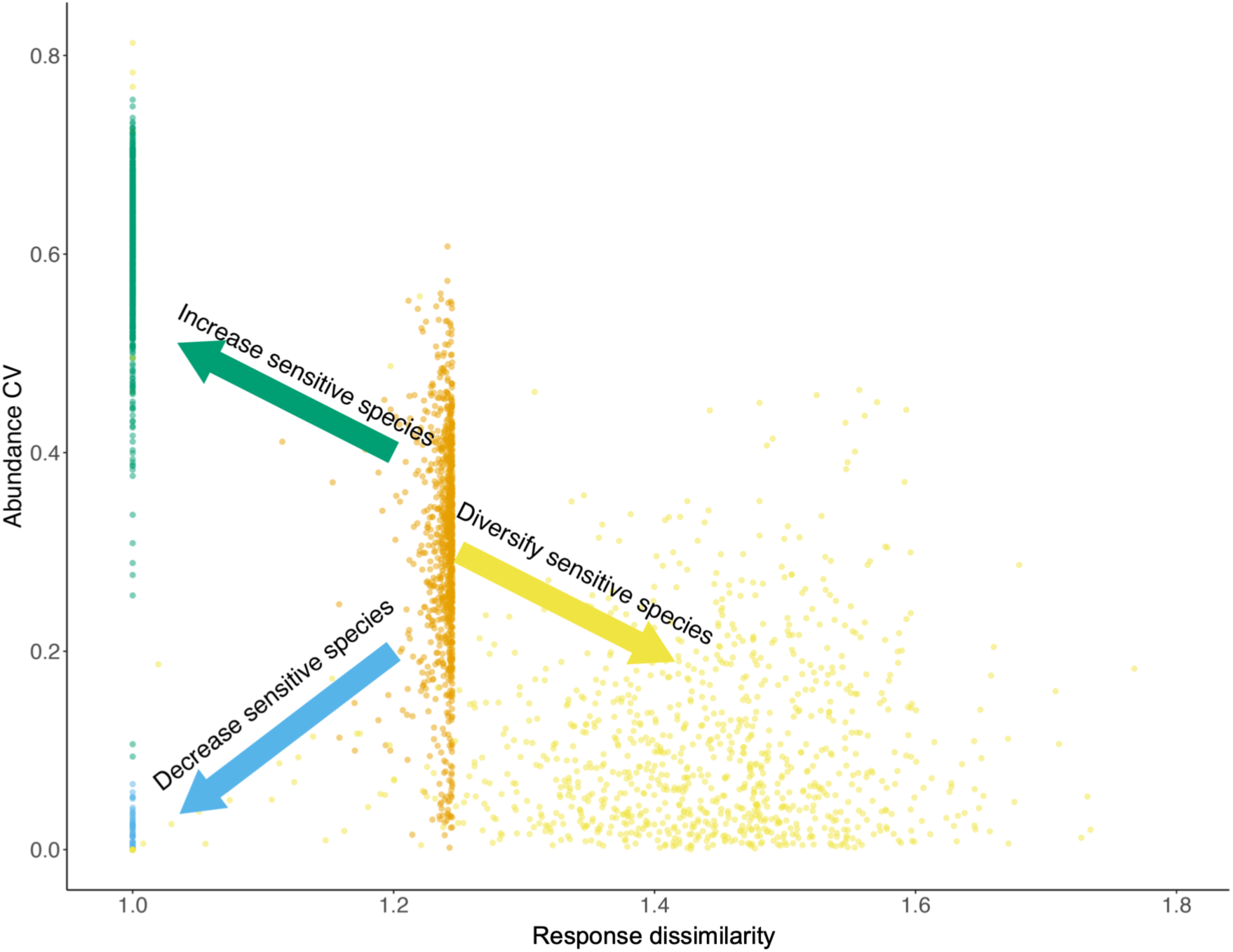
Response diversity can either stabilize or destabilize community dynamics depending on how it is generated. The relationship between response dissimilarity and abundance CV is shown for simulations with connectance fixed at 0.5. Dot colors indicate different simulation scenarios. Blue dots represent Scenario 1, in which no species responded to environmental change. Orange dots represent Scenario 2 with 50 sensitive species that responded identically to the environment. Yellow dots represent Scenario 3 with 50 sensitive species that differed in their environmental response slopes. Green dots represent Scenario 2 with all 100 species sensitive and responding identically to the environment.

## 4 Discussion

Understanding how biodiversity promotes ecological stability remains a central challenge in ecology. Response diversity has increasingly been recognized as a key mechanism linking biodiversity and ecological stability (Ives et al. 1999, Yachi and Loreau 1999, Hooper et al. 2005, Winfree and Kremen 2009, Bartomeus et al. 2013, Sasaki et al. 2019, Ross et al. 2023, Danet et al. 2025). Although previous studies have emphasized differences in species’ environmental optima as key drivers of ecological stability (Sasaki et al. 2019, Muraina et al. 2021, Schnabel et al. 2021, White et al. 2023) and Ross et al. (2023) proposed a framework for quantifying response diversity, our simulations reveal that one simple but overlooked component of response diversity—the number of species sensitive to environmental change—can substantially alter the relationship between response diversity and ecological stability. Insensitive species may become common when environmental changes are small in amplitude, when species differ in niche breadth as well as in their environmental optima, or when species exhibit physiological or phenotypic buffering. Specifically, we found that response dissimilarity, realized species richness, and community instability (abundance CV) varied with the number of sensitive species and variation in their responses. Although these factors had much smaller effects on realized species richness than connectance (May 1972), they had strong and sometimes counterintuitive effects on response dissimilarity, abundance CV, and the relationship between them.

In the absence of variation among sensitive species (Scenario 2), abundance CV increased monotonically with the number of sensitive species (Fig. 2B). This pattern likely arose because sensitive species fluctuated synchronously in response to environmental variation, thereby amplifying fluctuations in total community abundance. Although the influence of the number of sensitive species has largely been overlooked, this pattern is consistent with the predictions of Yachi and Loreau (1999). In their model, the CV of ecosystem productivity is greater when all species fluctuate in perfect synchrony (analogous to *M* = 100 in our Scenario 2) than when species exhibit imperfect synchrony (analogous to *M* < 100). These results suggest that species interactions played a smaller role in generating this pattern because connectance had much weaker effects on abundance CV than the number of sensitive species in our simulations, and the model of Yachi and Loreau (1999) assumed zero connectance. Notably, response dissimilarity showed a unimodal relationship with the number of sensitive species (Fig. 2C), reaching its maximum when half of the species were sensitive and the other half were insensitive. Consequently, the relationship between response dissimilarity and abundance CV was non-monotonic (Figs. 4 and S3): abundance CV increased with response dissimilarity when the number of sensitive species was less than *N/*2, but decreased with response dissimilarity when it exceeded *N*/2. Therefore, ignoring the number of sensitive species can obscure the relationship between response diversity and ecological stability.

Quantitative variation among sensitive species altered how the number of sensitive species affected response dissimilarity and abundance CV (Scenario 3, Fig. 3). First, response dissimilarity increased monotonically with the number of sensitive species and was generally higher than in Scenario 2, which assumed binary variation in species’ responses (compare Figs. 2C and 3C). This result is expected because sensitive species differed in their environmental responses, whereas insensitive species showed no response; consequently, increasing the number of sensitive species increased variation in environmental responses across the community. Second, quantitative variation among sensitive species reduced the effect of the number of sensitive species on abundance CV (compare Tables S3 and S4), although abundance CV and response dissimilarity remained positively associated (Fig. S4). In other words, whereas increasing the number of sensitive species destabilized communities, variation among sensitive species weakened this destabilizing effect. These findings suggest that the stabilizing effect commonly attributed to response diversity through species asynchrony (Sasaki et al. 2019, Muraina et al. 2021, Schnabel et al. 2021, White et al. 2023) depends on how that asynchrony is generated. For example, if species asynchrony arises because more species respond to the focal environmental variable, response dissimilarity increases but community instability may also increase. In contrast, when species asynchrony increases without changing the number of sensitive species, greater response diversity is associated with lower community instability, consistent with previous studies (Sasaki et al. 2019, Muraina et al. 2021, Schnabel et al. 2021).

Overall, our simulations indicate that biodiversity can influence ecosystem stability not only through changes in response diversity itself but also through changes in the composition of sensitive and insensitive species that generate response diversity. More specifically, the mechanisms generating response diversity can alter the observed relationship between response diversity and community stability (Fig. 4). When response diversity increased through greater variation among sensitive species (orange and yellow dots in Fig. 4), abundance CV was negatively associated with response dissimilarity. In contrast, when response diversity was generated by changes in the number of sensitive species (blue and orange dots in Fig. 4), abundance CV was positively associated with response dissimilarity (see also Fig. S3). Thus, response dissimilarity alone cannot distinguish between changes in the number of sensitive species and changes in variation among their responses, thereby allowing both positive and negative relationships between response diversity and community stability to emerge when the proportion of sensitive species is ignored. Our results provide one possible explanation for the weak relationship between response dissimilarity and ecological stability reported by Ross et al. (2023). Quantifying the numbers of sensitive and insensitive species, in addition to response diversity itself, will improve our understanding of biodiversity–stability relationships.

Although we used a generalized Lotka–Volterra (gLV) model to investigate the effects of response diversity, such models may not fully capture community dynamics arising from explicit interaction mechanisms (Momeni et al. 2017). Previous studies have identified conditions under which gLV models provide appropriate descriptions of community dynamics (Dedrick et al. 2023, Picot et al. 2023), for example in communities maintained under constant resource inflow and outflow. Here, we assumed that these conditions were satisfied and focused on the phenomenological relationship between response diversity and community stability without explicitly modeling the mechanisms underlying species interactions. An important next step will be to examine whether our conclusions hold in consumer–resource models and other mechanistic frameworks. Such models would also allow researchers to investigate situations in which environmental change affects not only intrinsic growth rates but also interspecific interactions, thereby revealing whether the importance of the number of sensitive species extends beyond phenomenological community models.

Another important direction for future research is to investigate the conditions under which different response diversity metrics are most informative. Although Ross et al. (2023) proposed two response diversity metrics—dissimilarity and divergence—we focused on response dissimilarity because response divergence showed little variation in our simulations (Fig. S1). In contrast, response divergence was informative in a recent empirical study (Polazzo et al. 2024). This difference likely reflects differences in model assumptions and data structure. In our model, response divergence was always zero in Scenarios 1 and 2. Although Scenario 3 allowed non-zero divergence, the metric was typically close to one whenever it differed from zero. This pattern likely resulted from our assumption that the distribution of species responses (*S_i_*) was symmetric (Eq. 6). More generally, different response diversity metrics may capture different aspects of species responses, and the distribution of those responses may determine which metric is most appropriate for a given ecological system.

In summary, this study provides a theoretical baseline for understanding how response diversity relates to species richness and community stability through a simple mathematical model. We showed that when response diversity is generated solely by changes in the number of environmentally sensitive species, the relationship between response diversity and community stability can be either positive or negative. These findings demonstrate that current response diversity metrics do not explicitly capture information about the number of sensitive species. Our results highlight that not only the magnitude of response diversity but also the biological processes generating it should be considered when evaluating biodiversity–stability relationships. We therefore suggest that future empirical and theoretical studies explicitly quantify the number of sensitive species and develop response diversity metrics that distinguish species sensitivity from variation in responses among sensitive species. These two components represent distinct ecological processes with potentially contrasting consequences for community stability.

## Acknowledgments

We thank members of Yamamichi laboratory and Toju laboratory for their helpful comments. We used ChatGPT for English proofreading.

## Author contributions

Conceptualization: SS, MY, HF, HT

Formal analysis, Investigation, Visualization, Writing - Original Draft Preparation: SS

Supervision: MY

Wiring- Review & Editing: SS, MY, HF, HT

Funding Acquisition: MY, HT

## Data availability statement

Data and codes are available on Zenodo (DOI: 10.5281/zenodo.21817794).

## Funding

This study was supported by Japan Science and Technology Agency (JST) Core Research for Evolutional Science and Technology (CREST) JPMJCR23N5.

## Competing interests

HT is a founder and director of Sunlit Seedlings Ltd. Other authors declare no competing interests.

## Notes

https://doi.org/10.5281/zenodo.21817794

