## Supplementary Materials for "Response diversity can stabilize or destabilize community dynamics depending on the number of insensitive species"

### Contents

Table S1. Quantile logistic regression analysis of realized species richness in Scenario 2. 2

Table S2. Quantile linear regression analysis of abundance CV in Scenario 2. 3

Table S3. Quantile logistic regression analysis of realized species richness in Scenario 3. 4

Table S4. Quantile linear regression analysis of abundance CV in Scenario 3. 5

Figure S1. Distributions of response divergence and response dissimilarity. 6

Figure S2. Realized species richness and abundance CV in Scenario 1. 7

Figure S3. Relationship between response dissimilarity and abundance CV in Scenario 2. 8

Figure S4. Relationship between response dissimilarity and abundance CV in Scenario 3. 9

**Table S1. Quantile logistic regression analysis of realized species richness in Scenario 2.**

| Variable | Connectance × Sensitive species | Connectance × Response dissimilarity |
|---|---:|---:|
| Intercept | 4.703 | 4.545 |
| Connectance | −12.164 | −13.005 |
| Sensitive species | −3.865 | |
| Response dissimilarity | | −3.513 |
| Connectance × Sensitive species | 5.167023 | |
| Connectance × Response dissimilarity | | 11.106 |
| AIC | 249377 | 245697 |

Blank cells represent the absence of the variable in the regression model. All variables were scaled between 0 and 1.

**Table S2. Quantile linear regression analysis of abundance CV in Scenario 2.**

| Variable | Connectance × Sensitive species | Connectance × Response dissimilarity |
|---|---:|---:|
| Intercept | 0 | 0 |
| Connectance | 0 | 0 |
| Sensitive species | 0.55 | |
| Response dissimilarity | | 0.213 |
| Connectance × Sensitive species | 0.078 | |
| Connectance × Response dissimilarity | | 0.067 |
| AIC | −129147.7 | −36957.87 |

Blank cells represent the absence of the variable in the regression model. All variables were scaled between 0 and 1.

**Table S3. Quantile logistic regression analysis of realized species richness in Scenario 3.**

| Variable | Connectance × Sensitive species | Connectance × Response dissimilarity |
|---|---:|---:|
| Intercept | 4.669 | 4.337 |
| Connectance | −12.054 | −11.681 |
| Sensitive species | −4.897 | |
| Response dissimilarity | | −3.109 |
| Connectance × Sensitive species | 11.478 | |
| Connectance × Response dissimilarity | | 8.582 |
| AIC | 159262.8 | 163252.6 |

Blank cells represent the absence of the variable in the regression model. All variables were scaled between 0 and 1.

**Table S4. Quantile linear regression analysis of abundance CV in Scenario 3.**

| Variable | Connectance × Sensitive species | Connectance × Response dissimilarity |
|---|---|---|
| Intercept | 0.002 | 0.002 |
| Connectance | −0.003 | −0.002 |
| Sensitive species | 0.058 | |
| Response dissimilarity | | 0.072 |
| Connectance × Sensitive species | 0.14 | |
| Connectance × Response dissimilarity | | 0.181 |
| AIC | −126337.3 | −128418.4 |

Blank cells represent the absence of the variable in the regression model. All variables were scaled between 0 and 1.

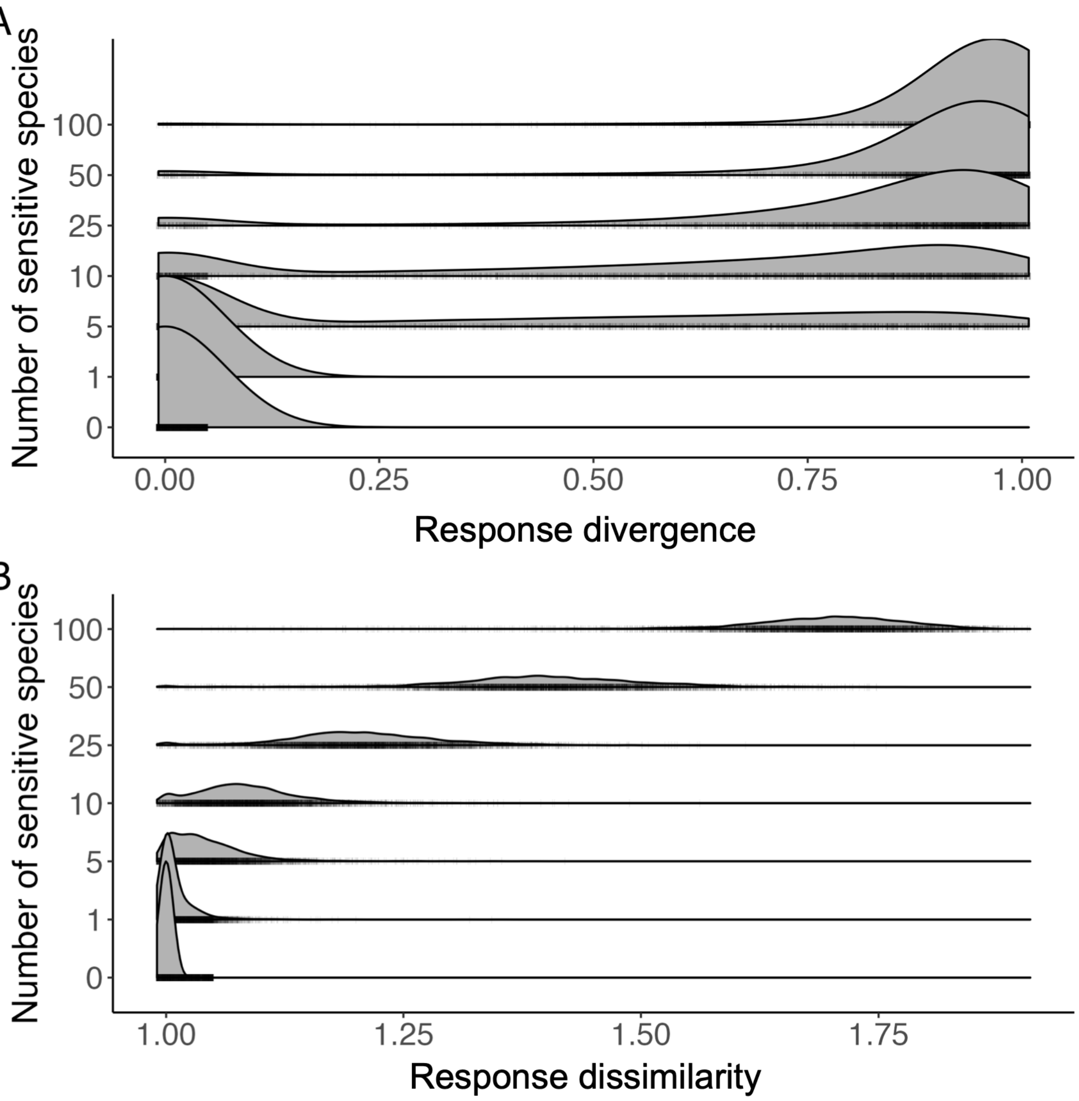


**Figure S1. Distributions of response divergence and response dissimilarity.** Response divergence (A) and response dissimilarity (B) were calculated from simulations in Scenario 3 after excluding simulations in which realized species richness was zero or one. Vertical black lines indicate the values of the two metrics obtained from individual simulations.

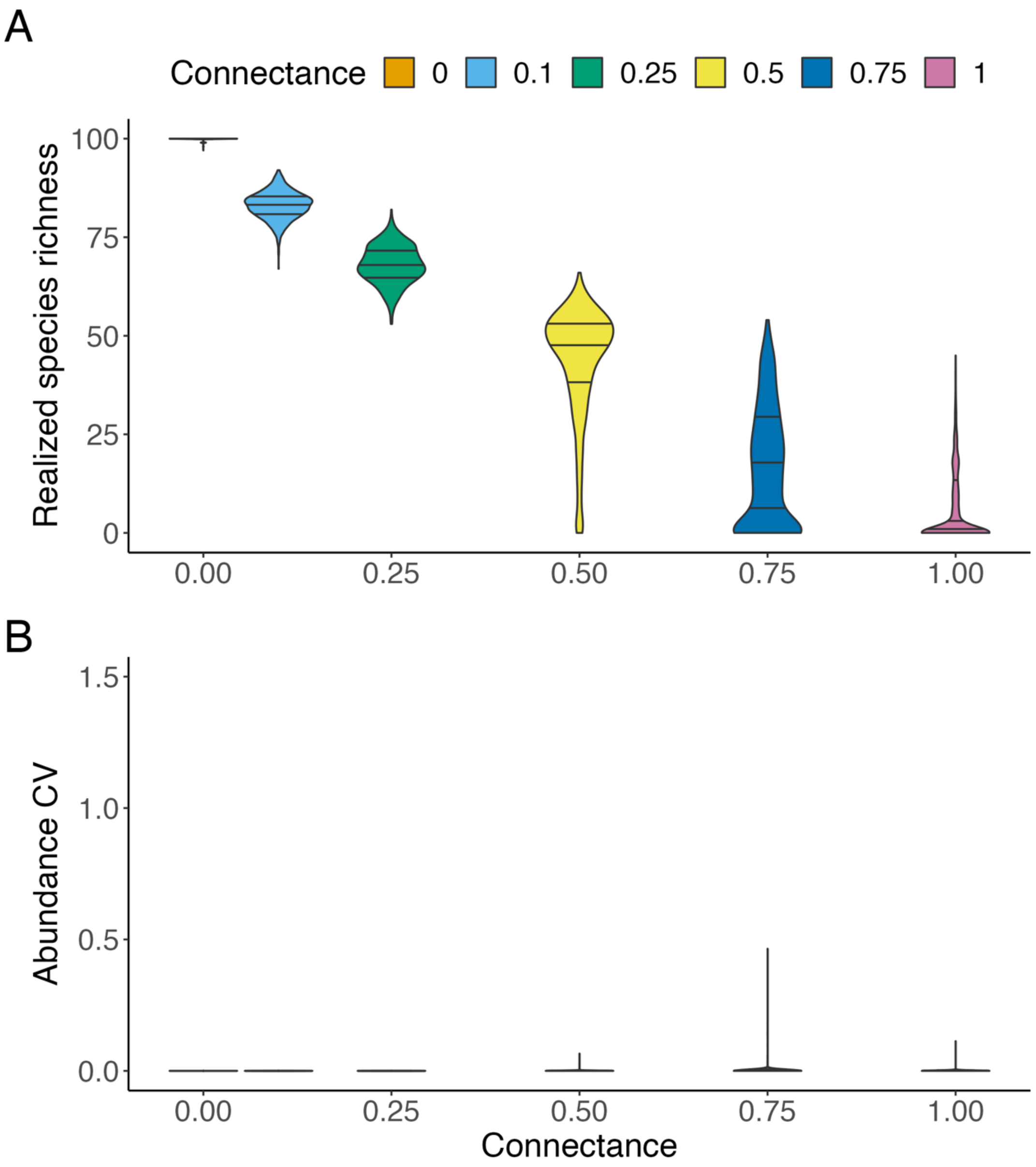


**Figure S2. Realized species richness and abundance CV in Scenario 1.** Realized species richness (A) and abundance CV (the coefficient of variation of total community abundance calculated over the interval from $t$ = 4,900 to $t$ = 5,000) (B) are shown. Horizontal lines indicate the fitted quantile regression lines. These results correspond to the leftmost columns of Figs. 2 and 3, where the number of sensitive species was zero.

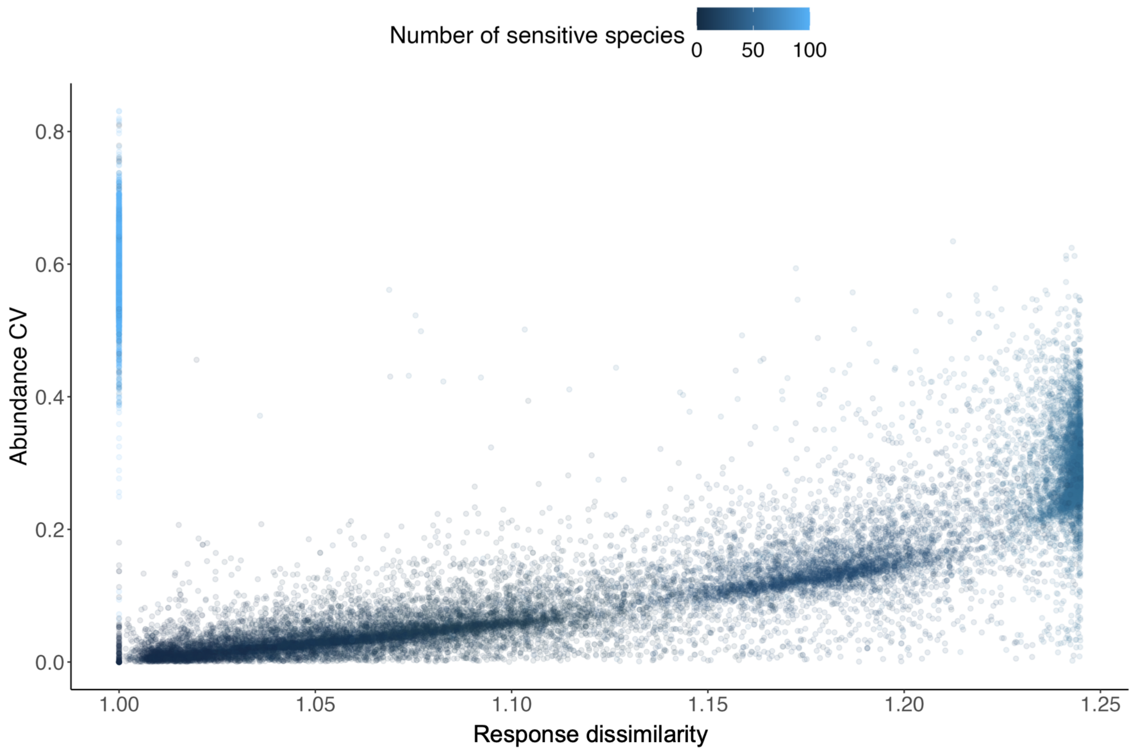


**Figure S3. Relationship between response dissimilarity and abundance CV in Scenario 2.** Dot color indicates the number of sensitive species among the $N = 100$ species. This figure presents the same data as Figs. 2B and 2C. Response dissimilarity was highest when the number of sensitive species was intermediate, whereas low response dissimilarity was observed when the number of sensitive species was either low or high. Consequently, low response dissimilarity was associated with either low or high abundance CV.

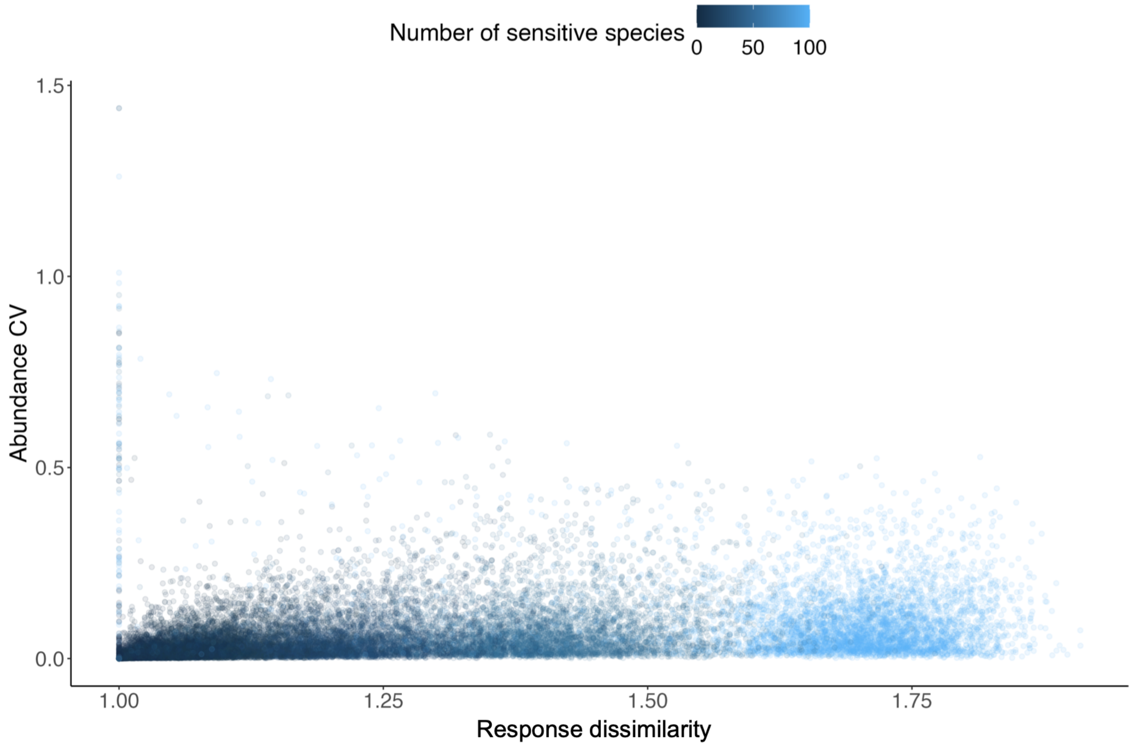


**Figure S4. Relationship between response dissimilarity and abundance CV in Scenario 3.** Dot color indicates the number of sensitive species among the $N = 100$ species. This figure presents the same data as Figs. 3B and 3C. Response dissimilarity generally increased with the number of sensitive species and was positively associated with abundance CV. The Pearson correlation coefficient between response dissimilarity and abundance CV was 0.473 (95% confidence interval: 0.465–0.480).
